# A bacterial tragedy of the commons that masks the actual frequency of mutants

**DOI:** 10.1101/2020.04.02.022889

**Authors:** Henrique Iglesias Neves, Gabriella Trombini Machado, Taíssa Cristina dos Santos Ramos, Hyun Mo Yang, Ezra Yagil, Beny Spira

## Abstract

The frequency of mutants in a population is central to the understanding of evolution. Mutant frequency is usually assessed by plating a bacterial culture on selective medium in which only specific rare mutants can grow, assuming that all mutant cells present on the plate are able to form colonies. Here we show an exception to this rule. Wild-type *Escherichia coli* cells are unable to grow with glycerol-2-phosphate (G2P) as a carbon source. In contrast, PHO-constitutive mutants can hydrolyse G2P to glycerol and form colonies on plates having G2P as their sole carbon source. However, the frequency of PHO-constitutive colonies on the selective plate is exceptionally low. Here we show that such mutations occur at a relatively high rate, but the growth of the existing mutants is inhibited due to a competition with the surrounding wild-type cells for the limited amounts of glycerol produced by the mutants. This scenario in which neither the wild-type nor the majority of the mutants are able to grow constitutes an unavoidable case of the ‘tragedy of the commons’. Evidence shows that the few mutants that do form colonies derive from micro-clusters of mutants on the selective plate. In addition, a mathematical model describes the fate of the wild-type and mutant populations on the selective plate.

## Introduction

The rate at which mutations occur is characteristic of the species and is also influenced by the environment [1, 2, 3, 4]. Under stressful conditions, a fraction or the entire bacterial population increase the mutation rate by some amount in what appears to be a response mechanism to environmental challenges [1, 4, 5, 6].

Ever since Luria & Delbrück seminal paper [7], it became apparent that changes in the bacterial genome are not directed to a relevant locus, notwithstanding selective pressures or environmental condition. However, the idea of directed mutagenesis resurfaced when non-lethal selective conditions instead of selection to virus/antibiotic resistance were employed, a process that has been named "adaptive or directed mutagenesis" [8]. Later, Cairns and others have shown that some mutations arise in response to environmental challenges [4, 5, 8, 9, 10, 11]. Adaptive mutations must occur in non-dividing bacteria and only after the bacteria have been in contact with the selective medium. The very existence of adaptive mutations is still disputed and led to a controversy that is still unresolved [12, 13, 14, 15, 16].

Another factor that may affect the assessment of mutation rate/mutant frequency is the density of the bacterial population plated on the selective medium [17, 18, 19, 20]. On one hand, traces of usable nutrients may foment bacterial replication on the plate and falsely increase the number of mutant colonies [19, 20]. This especially occurs in mutational systems in which the selective plates are incubated for longer periods of time, usually more than 48 h, during which the bacterial population grow slowly using non-selective alternative nutrient traces. For this reason, it is a usual practice to plate “scavenger” or “filler” cells that cannot mutate to prototrophy, but can scavenge nutrient traces from the medium together with the test bacteria [17, 18, 19, 21]. On the other hand, *E. coli* grown to high densities have consistently shown lower rifampicin resistance mutation rates, in a mechanism dependent on cell–cell signaling and on the quorum sensing gene *luxS* [22]. This finding was recently extended to the yeast *Saccharomyces cereviseae*, in which an inverse relation between cell density and mutation rate has been also observed [23]. Hence, the use of scavenger cells in the experimental assessment of mutation rate in non-lethal selection systems such as the ability to use a nutrient source can lead to an underestimation of mutation rate due to density-associated mutation inhibition.

Genes belonging to the PHO regulon in *E. coli* are induced under conditions of or- thophosphate (Pi)-deprivation. The PHO regulon genes are controlled by the two-component system PhoB/PhoR. The histidine kinase PhoR receives a signal from the Pst (Pi-specific transport) system about the availability of Pi in the periplasm. Pst is encoded by *pstSCAB-phoU* operon, or in short *pst* operon. When Pi concentration declines below 4 μM, PhoR auto-phosphorylates and immediately transfers its Pi moiety to PhoB, which in turn binds to the PHO-box sequences in the promoter regions of all PHO genes thereby initiating the transcription of the PHO regulon genes [24]. Mutations in any one of the five *pst* operon genes result in the constitutive expression of the PHO regulon. i.e., in these mutants PhoR continuously phosphorylates PhoB keeping the expression of the PHO regulon genes at its highest even under conditions of excess Pi [25, 26]. Thus the Pst system, besides being the principal Pi-uptake system in *E. coli*, also acts as a transcriptional repressor of the PHO regulon [27, 28, 29].

PHO-constitutive *pst* mutants can be selected by plating wild-type bacteria on a minimal medium containing glycerol-2-phosphate (G2P) as the sole carbon (C) source [25]. Since *E. coli* cannot transport G2P, at least not in sufficient amounts to provide the C needed for growth, G2P must first be cleaved in the periplasm by the product of another PHO regulon member – *phoA*, that encodes alkaline phosphatase (AP). AP hydrolyzes G2P, releasing glycerol that enters the bacterial cell mostly by facilitated diffusion via GlpF [30, 31]. Thus, under high-Pi conditions the repressed level of AP activity in the periplasm is too low to provide sufficient glycerol for cell growth. Only PHO-constitutive mutants (PCMs), which carry a null mutation in one of the five *pst* operon genes [24] express high levels of AP that enable them to grow on G2P medium [26, 32]. Some mutations in *phoR* also result in the constitutive expression of the PHO regulon and likewise confer on the ability to grow on G2P. However, all PHO-constitutive mutations in strain MG1655 hitherto analyzed are in the *pst* operon [26]. We have previously shown that when PCMs are selected on G2P as a sole C source very few mutants emerge and this is despite the fact that the target for PHO-constitutive mutations is quite large (the entire *pst* operon plus a few bases in *phoR*). In addition, the first PCM colonies appear only on the third day of incubation, and their number on the selective plates increase with time in a sigmoid pattern. Having confirmed that PCMs, once isolated, form colonies on G2P plates within less than 48 hours, we suggested that the late emergence of PCMs on the selective medium could be explained by stress-induced adaptive mutagenesis [26]. However, further investigation reported here have demonstrated that the vast majority of PCMs in a wild type population are cryptic and that the actual spontaneous rate of the PCMs is in fact quite high. This conclusion was strengthened by finding out that PCMs growth on the selective plate is inhibited by the high density of wild-type bacteria in a competition for the scarce glycerol molecules released by the PCMs, resulting in a “tragedy of the commons” case [33, 34, 35] in which neither the majority of PCMs nor the wild-type cells are able to grow and form colonies.

## Results

### Characterization of the emerging PHO-constitutive mutants

The pattern of emergence of PCMs derived from the prototype *E. coli* K-12 strain MG1655 was investigated. In a typical experiment, an overnight culture containing 10^9^ wild-type MG1655 cells is plated on TG2PP-minimal medium containing G2P as the sole carbon source, excess Pi and the AP substrate XP that stains blue the constitutive mutants owing to their high AP activity. Figure 1a shows the kinetics of PCMs accumulation in 17 independent cultures. In all of them the first few colonies emerged on the 3^rd^ −4^th^ day of incubation, while more colonies kept appearing for the next 10 days, with the bulk of colonies emerging between the 5^th^ and the 8^th^ day following plating. The mean number of mutant colonies at the end of 16 days was 41.1 ± 17.9. Typically the mutant frequency oscillates between 20-200 mutants/plate (0.2 – 2.0 × 10^−7^ mutants/cell/generation). The distribution of mutants emergence per day is shown in Figure 1b. The general mutant frequency in *E. coli* is between 10^−6^ and 10^−7^ per gene [36, 37, 38]. Thus the expected mutant frequency for mutations in the five *pst* operon genes should have been 5 × 10^−7^ to 5 × 10^−6^/cell/generation, roughly 25–250 times higher than what has been observed here.

**Figure 1.**
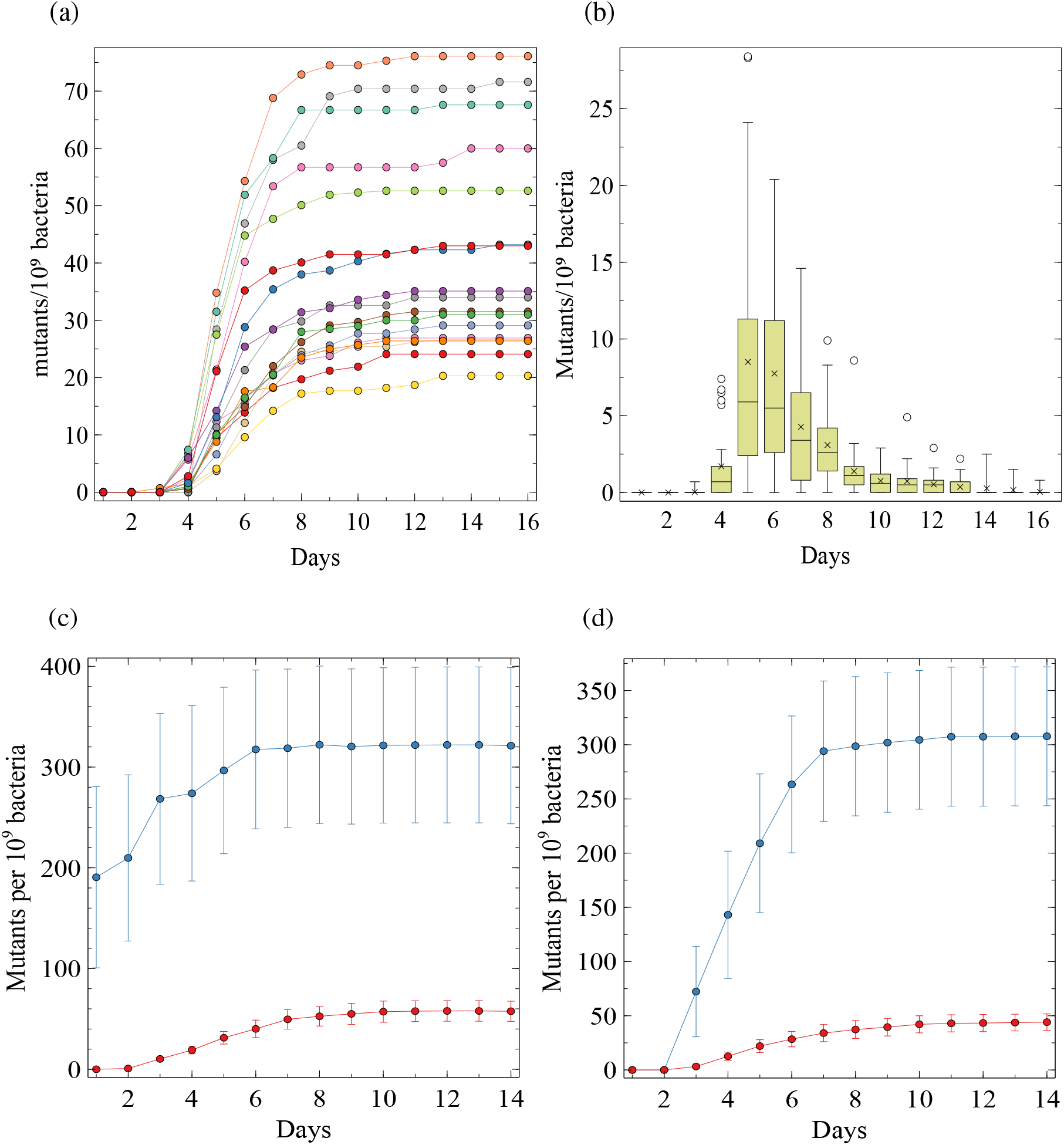
(a) Daily accumulation and distribution of PCMs in 17 independent cultures. 10^9^ MG1655 bacteria were plated on each TG2PP plate. Number of mutants emerging on the selective plates recorded for 16 days. (b) Boxplots representing the distribution of new colonies appearing on each day in the 17 plates. (x), (-) and (○) represent the means, medians and outliers, respectively. (c) Frequency of PCMs and rifampicin resistant mutants. 10^9^ bacteria (MG1655) were plated on TGP minimal medium supplemented with rifampicin (100 μg/ml) or on TG2PP plates. 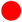, PCMs; 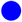, Rif^R^ mutants. Each point represents the mean ± S.E.M. of 10 independent cultures. (d) Selection of PCMs in a *mutS* background. 10^9^ MG1655 or !:: *mutS*::Cm bacteria were each separately plated on TG2PP plates. 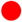, PCMs from MG1655; 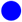, PCMs from *mutS*. Each point represents the mean ± S.E.M. of seven in-dependent cultures.

To test whether the late emergence of PCM colonies and their low mutant frequency is specific to PCMs selection, the frequency of rifampicin-resistant mutants (Rif^R^) was compared. Bacteria were grown overnight in medium TGP and plated on TGP containing rifampicin (100 μg/ml) and on TG2PP plates (Figure 1c). The frequency of Rif^R^ mutants at the 14^th^ day was 3.2 × 10^−7^/*generation*, which is compatible with data reported elsewhere for strain MG1655 [39, 40], but different from others that reported lower frequencies [41, 42, 43]. In contrast, bacteria plated on TG2PP showed the usual pattern of mutant accumulation (first visible colonies at day 3 and a low mutant frequency of 5.8 × 10^−8^/generation, calculated at the 14^th^ day). Fifty nine percent of the Rif^R^ mutants were already apparent on their selective plates within 24 hours, 83% on day 3 and 98% on day 6. The high frequency of Rif^R^ colonies that were evident in the first 3 days indicate preexisting spontaneous mutations in the overnight cultures. By comparison, on average only 17% of the PCM colonies emerged on the selective plates at day 3 and more of them kept appearing up to day 10. It is worth mentioning that upon restreaking on G2P plates even the late PCMs form colonies within 48 h. We also tested the pattern of emergence of PCMs in a Δ*mutS* derivative of strain MG1655 (strain RI103). The Δ*mutS* strain is partially deficient in DNA mismatch repair and displays a 10-100 fold increase in mutation rate [44, 45, 46]. Figure 1d shows a 7.5 fold increase in the mean number of PCMs derived from the *mutS* strain at the 14^th^ day. However, not a single colony emerged before the 3^rd^ day and the pattern of mutant accumulation was similar to the one observed in the wild-type strain, suggesting that the late emergence of PCMs is not necessarily associated with the low frequency of these mutants.

The emergence of PCM colonies appears as adaptive in nature, i.e., the mutants do not emerge at the early days (pre-existent mutations), but they appear only after the bacteria are exposed to the selective conditions (G2P as the C source). However, in other adaptive mutational systems, such as the classical *lacZ* reversion assay, known as the Cairns system [8, 17], in addition to the late adaptive revertants pre-existent mutants do appear in 48 h with a frequency that is higher than 10^−7^/bacteria/generation. Furthermore, the emergence of adaptive *lacZ*^+^ revertants follow a Poisson distribution. In contrast, the low frequency of PCMs and the large variance across different cultures of the same strain (Figure 1a mean and variance at the end of the experiment were 41 and 320, respectively) follows a Luria-Delbrück distribution, and suggests that PHO-constitutive mutations are not adaptive in nature. Thus, the apparent absence of pre-existent PCMs and their low frequency may imply another phenomenon such as growth inhibition of these mutants on the selective plate. The following experiments were aimed at investigating this possibility.

One hundred Δ*pst*∷Km bacteria (strain BS07) were mixed with increasing numbers (10^5^, 10^6^, 10^7^, 10^8^ or 10^9^) of MG1655 wild-type cells and plated on TG2PP. The emergence of PCMs was examined after 48 hours (Figure 2). When 0 or 10^5^ wild-type bacteria were plated together with the Δ*pst*∷Km cells, virtually all Δ*pst* colonies were recovered. However, higher concentrations of wild-type cells progressively inhibited the emergence of Δ*pst* colonies. The inhibition levels were on average 47%, 86%, 99% and 100% for mixes containing 10^6^, 10^7^, 10^8^ and 10^9^ wild-type cells, respectively. This result indicates that as of a certain density the neighboring wild-type cells inhibited the growth of the PCM.

**Figure 2.**
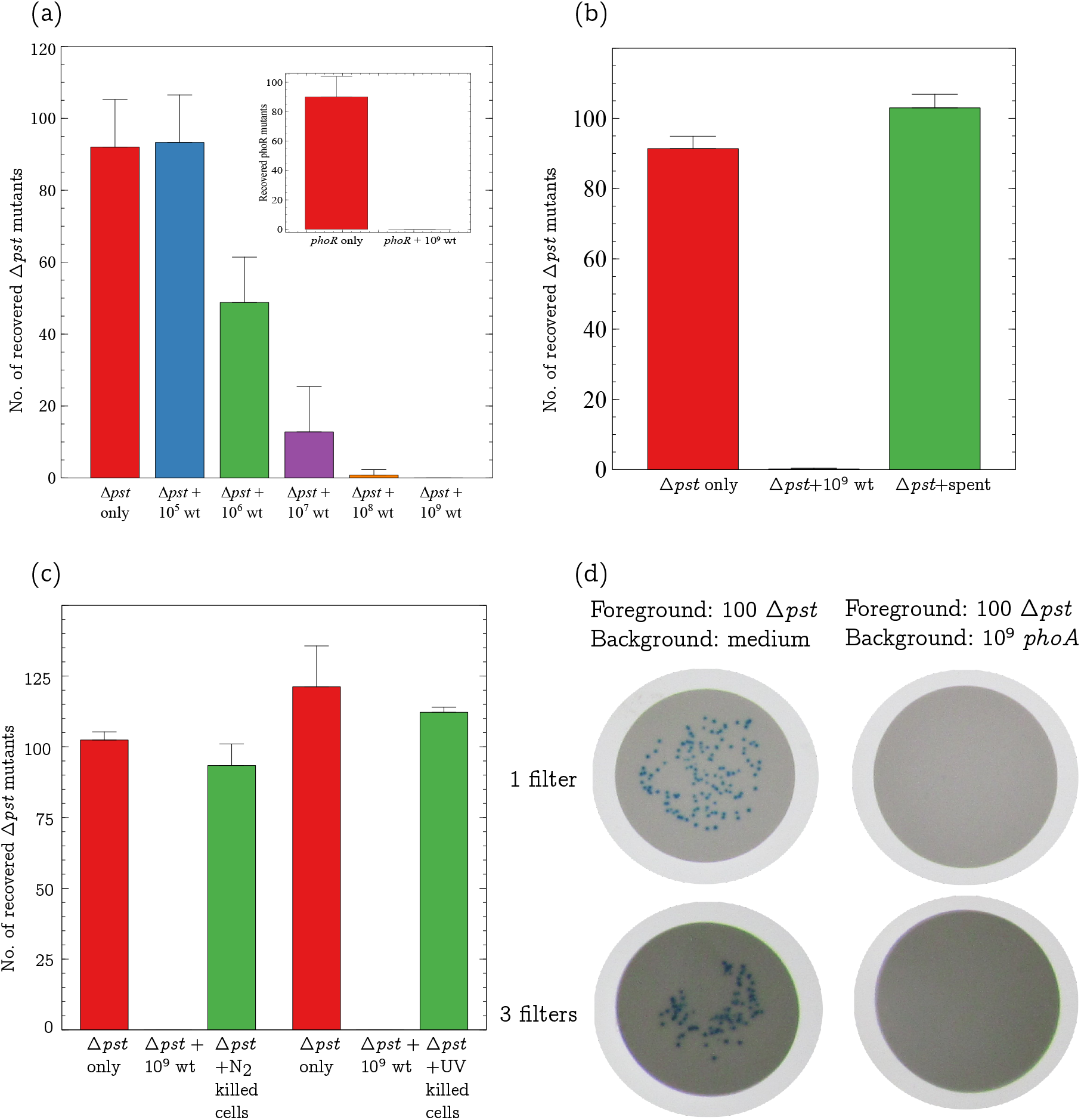
**(a)** Growth inhibition of Δ*pst*∷Km cells plated with increasing amounts of wild-type MG1655 bacteria (wt). Inset shows the growth of *phoR* cells plated with and without 10^9^ wild-type cells. Each bar represents the mean ± S.E.M. of 5 independent experiments. **(b)** Effect of wild-type spent on PCMs growth. 10^9^ MG1655 cells grown overnight in medium TGP, were resuspended in TG2PP and further incubated at 37°C for 48 h. The filtered sup of this culture (spent) was mixed with 100 Δ*pst*∷Km cells and plated on TG2PP for another 48 hours (Δ*pst* + spent). 100 Δ*pst*∷Km (Δ*pst* only) and 100 Δ*pst*∷Km mixed with 10^9^ wild-type cells (Δ*pst* + 10^9^ wt) were plated as controls. **(c)** Growth inhibition by dead cells. 10^9^ wild-type bacteria were killed either by freeze-thawing in liquid nitrogen (N_2_ killed cells) or by UV-irradiation (UV killed cells). The dead bacteria were mixed with 100 Δ*pst* cells, plated on TG2PP and incubated for 48 h. Δ*pst* cells alone and Δ*pst* cells mixed with 10^9^ untreated wild-type cells served as controls. Each bar represents the mean ± S.E.M. of 5 independent cultures. **(d)** Growth inhibition of Δ*pst* PCMs does not require contact with inhibitor cells. Δ*pst*∷Km cells were grown overnight in medium TGP, washed and diluted in saline. Approximately 100 bacteria were spread on the surface of a 0.22 μm filter which in turn was placed on a TG2PP plate seeded (left) or unseeded (right) with 10^9^ Δ*phoA*∷Cm cells. In addition, 100 Δ*pst*∷Km bacteria were spread on top of a 3 filter stack placed on the surface of a TG2PP plate. The plates were incubated at 37°C for 48 h.

To test whether this inhibitory effect likewise acts on another PCM genotype, 100 cells of a *phoR* mutant (strain RI65), which is also PHO-constitutive, were mixed with 10^9^ wild-type bacteria and plated on TG2PP medium. The inset in Figure 2 shows that the *phoR* mutants were likewise completely inhibited by the presence of the wild-type cells.

### Mutant frequency and mutation rate of PHO-constitutive mutants

The growth inhibition of PCM colonies by wild-type cells may mask the real frequency of these mutants on the selective plate and disguise their mutation rate. To bypass this inhibition and to assess the rate of PCMs a Luria-Delbrück fluctuation test with a limited amount of bacteria was conducted. Approximately 1000 MG1655 bacteria were inoculated in each of 60 tubes containing 0.1 ml of medium TGP supplemented with only 110 nM glucose and grown for 30 hours. This low glucose concentration limits cell yield to ~ 5 · 10^4^ bacteria/ml, which is below the threshold that inhibits the propagation of PCM cells (Figure 2a). The entire volume of each culture was then plated on 60 TG2PP plates and the number of colonies formed at 48 hours was counted (Table 1). Of the 60 plates only 6 showed some PCM colonies. The mutation rate of PCM, calculated by the Luria-Delbrück P_0_ method [7, 47], was 7.9 × 10^−6^ mutations/cell/generation. The same experiment was repeated using the mutator strain Δ*mutS*∷Cm (strain RI1103) in which case 57/60 plates showed PCM colonies. This result allowed us to calculate the mutation rate by additional estimators that rely on medians, such as the Lea-Coulson [48] and Jones method [49]. The calculated mutation rates in this strain were between 9.2 × 10^−5^ and 1.0 × 10^−4^ mutations/cell/generation, which is roughly 12 times higher than in the wild type strain. Thus the estimated frequency of PCMs among 10^9^ wild-type bacteria seeded on one TG2PP plate should be at least 8, 000, but less than 100 colonies are usually observed at the end of 14 days (Figure 1). In the case of the *mutS* strain, an even higher number of mutations (~ 96, 000) would be expected.

**Table 1.** Fluctuation test for PCMs in strains MG1655 and Δ*mutS*.

| | MG1655 | $\Delta mutS$ |
| --- | --- | --- |
| **Plates with 0 mutants** | 54 | 3 |
| **Total number of cultures** | 60 | 60 |
| $\mu$ (/cell/generation) ($P_0$)<sup>1</sup> | $7.9 \times 10^{-6}$ | $9.2 \times 10^{-5}$ |
| $\mu$ (/cell/generation) (Lea-Coulson)<sup>2</sup> | – | $1.01 \times 10^{-4}$ |
| $\mu$ (/cell/generation) (Jones)<sup>3</sup> | – | $9.2 \times 10^{-5}$ |

<sup>1</sup> Luria-Delbrück $P_0$ method [7]
<sup>2</sup> Lea & Coulson method [48]
<sup>3</sup> Jones estimator [49]

In addition to the fluctuation test, an alternative strategy was employed to estimate the actual number of plated PCM cells (mutant frequency) on the selective plate. Figure S1 displays a scheme representing the experimental design. Five overnight cultures each containing 10^9^ wild-type MG1655 cells were mixed with increasing numbers of Δ*pst*∷Cm cells (from 10^3^ to 10^5^) and plated on TG2PP (Table 2, 1^st^ column). After 7 days of incubation, PCM colonies from each mix were randomly chosen and replica plated on L-agar containing chloramphenicol to determine the number of Δ*pst*∷Cm mutants (3^rd^ column). Based on the proportion of Δ*pst*∷Cm inhibition, the number of spontaneous PCM colonies on the selective plate was calculated. Once the number of Δ*pst*∷Cm cells added is known, their recovery level reveals the extent of their inhibition (5^th^ column). From this, the expected number of the spontaneous PCM cells that were plated on the selective plates (6^th^ column) could be derived. For instance, of the 345 colonies that emerged on the 5 plates seeded with the mix of 10^9^ wild-type bacteria + 5,000 Δ*pst*∷Cm cells, only 26 proved to be Δ*pst*∷Cm, indicating an inhibition factor of 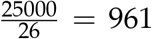 fold. Given that on the 5 plates there were in total only 319 spontaneous PCM, the real number of PCMs on these plate should have been 319 × 961 = 3.06 × 10^5^ per 5 × 10^9^ cells (6.1 × 10^4^/10^9^ cells). The last lane in Table 2 shows the mean inhibition factor was 897 and the expected number of PCMs on a plate seeded with a culture of 10^9^ MG1655 cells was 50,000 colonies. Thus the average expected frequency of PCMs in a TG2PP plate seeded with 10^9^ cells is 5.0 · 10^−5^, which is about 6.8 times above the estimated mutation rate of 7.9 × 10^−6^ obtained by the fluctuation test (Table 1). The ratio [mutant frequency]/[mutation rate] found here is similar to the observed in other systems [7, 50, 51].

**Table 2.** Inferring the frequency of spontaneous PCMs in a mixed culture.^1^.

| 5 plates each containing $10^9$ wt cells + following no. of $\Delta pst::Cm$ cells | Total PCMs at day 7 | $\Delta pst::Cm$ colonies | Spontaneous PCMs colonies | Inhibition factor ($\frac{\Delta pst::Cm \text{ plated}}{\Delta pst::Cm \text{ observed}}$) | Expected n° of spontaneous PCMs/$10^9$ cells |
| --- | --- | --- | --- | --- | --- |
| $(5 \times) 10^3$ | 267 | 7 | 260 | 714 | $3.7 \times 10^4$ |
| $(5 \times) 3 \times 10^3$ | 269 | 20 | 249 | 750 | $1.9 \times 10^4$ |
| $(5 \times) 4 \times 10^3$ | 364 | 18 | 346 | 1111 | $7.7 \times 10^4$ |
| $(5 \times) 5 \times 10^3$ | 345 | 26 | 319 | 961 | $6.1 \times 10^4$ |
| $(5 \times) 10^4$ | 209 | 31 | 178 | 1612 | $5.7 \times 10^4$ |
| $(5 \times) 2.5 \times 10^4$ | 254 | 114 | 140 | 658 | $1.8 \times 10^4$ |
| $(5 \times) 5 \times 10^4$ | 977 | 391 | 586 | 639 | $7.5 \times 10^4$ |
| $(5 \times) 10^5$ | 1098 | 686 | 412 | 729 | $6.0 \times 10^4$ |
| Means | | | | 897 | $5.0 \times 10^4$ |

<sup>1</sup> Five cultures each containing $10^9$ wild-type bacteria were mixed with increasing numbers of $\Delta pst::Cm$ mutants and immediately plated on TG2PP. After 7 days of incubation, the total number of colonies was counted and tested for chloramphenicol resistance. The ratio [ $\Delta pst$ plated]/[ $\Delta pst$ colonies observed] gave the inhibition factor of $\Delta pst$ . The expected n° of PCM/ $10^9$ cells was calculated by multiplying the number of spontaneous PCMs on the 5 plates by the inhibition factor divided by 5.

### Mechanism of PCMs growth inhibition

In quorum sensing bacteria send signals to their counterparts or other organisms through the secretion of small molecular weight molecules. In some cases, quorum sensing molecules mediate a signal for growth inhibition or cell death [52]. To test whether growth inhibition of PCMs is mediated by a secreted molecule, the filtered spent medium of an MG1655 culture grown overnight in TGP and further incubated in liquid TG2PP for 48 hours (to mimic the conditions of PCMs selection), was used to resuspend 100 Δ*pst* bacteria, which were then plated on TG2PP and incubated for 48 hours. Figure 2b shows that the supernatant of the wild-type strain did not inhibit the growth of the Δ*pst* colonies. We also tested whether the wild-type cells must be alive in order to inhibit the growth of PCM. 10^9^ wild-type bacteria killed either by immersion in liquid nitrogen or by UV-irradiation, were mixed with 100 Δ*pst* mutants and plated on TG2PP for 48 hours. Figure 2c shows that wild-type bacteria killed by either method failed to inhibit the growth of PCM colonies on the selective plate indicating that the inhibitor cells must be alive to inhibit PCMs.

Other instances of reported growth inhibition systems involve either cell contact [53, 54, 55] or proximity [56]. To test whether PCMs growth inhibition depends on contact between the mutant and wild-type bacteria, 100 Δ*pst* cells were spread on top of a 0.22 μm membrane placed over a TG2PP plate seeded with 10^9^ *phoA* cells and incubated for 48 h (Figure 2d). In the absence of 10^9^ bacteria on the plate, all 100 Δ*pst* cells placed on the filter formed PCM colonies within 48 h. However, when the filter was placed on a TG2PP plate previously seeded with the *phoA* cells, not a single colony emerged, even when the bacteria were placed on top of three filters. Thus, growth inhibition does not require contact between the PCMs and the surrounding bacteria. It should be noticed that the *phoA* mutant inhibits PCMs growth as does the wild-type strain (see Figure 3a and Figure 3c), but unlike the wild-type strain it cannot generate PCMs even after long periods of incubation in the presence of G2P due to the lack of AP [26].

**Figure 3.**
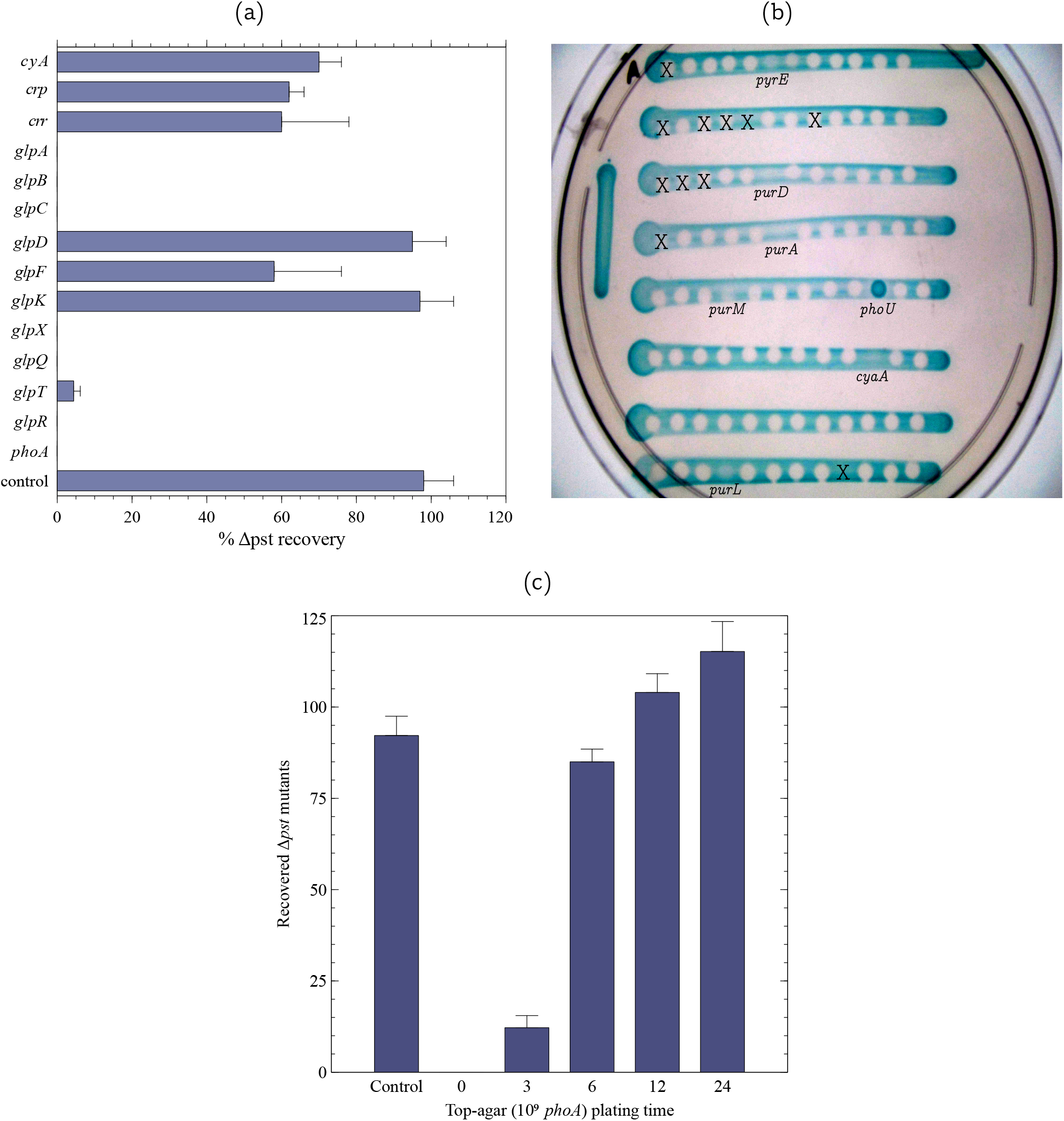
**(a)** Inhibition of Δ*pst* growth by single gene knockouts from Keio collection (BW25113 background). One hundred Δ*pst* cells were mixed with 10^9^ bacteria carrying individual deletions in each of the following genes: *cyaA, crp, crr, glpA, glpB, glpC, glpD, glpF, glpK, glpX, glpQ, glpT, glpR* and *phoA* as a positive control. The plates were incubated for 2-3 days at which time the PCM colonies were counted. ‘Control’ represents the Δ*pst* strain plated in the absence of other bacteria. Each bar represents the mean ± S.E.M. of at least 3 independent cultures. **(b)** Screening the *E. coli* collection of knockouts for Δ*pst* growth inhibition. Δ*pst* cells and the library knockouts (Keio collection plate nº 53) were grown overnight in TGP medium. The Δ*pst* culture was diluted a hundredfold and 30 μl of this dilution was used to create a linear patch on a TG2PP plate supplemented with XP. Once the Δ*pst* patch was dry, 2 μl of each knockout strain were dropped over the patches. The plates were incubated for 48 hours at 37°C. In the vast majority of cases a halo was formed inside the patch where the knockout strain was applied, indicating that Δ*pst* growth was inhibited by this particular strain. Only a few knockouts described in the main text allowed the growth of Δ*pst*, characterized by a bluish color inside the drop. These strains were further tested in a conventional inhibition assay, as in Figure 3a to confirm this phenotype. The strong blue color inside the *phoU* drop is because this mutant is a PCM that grows on TG2PP. Of those that did not inhibit Δ*pst*, the majority was formed by auxotrophic strains. For instance, in plate 53, the knockouts of *pyrE, purA, purD, purL* and *purM* did not inhibit Δ*pst* growth, but they do not grow or grow very poorly in minimal medium. Halos marked with an X correspond to bacteria that are not part of the Keio collection (see the Keio collection documentation at https://shigen.nig.ac.jp/ecoli/strain/resource/keioCollection/about). **(c)** Formation of PCM colonies from apparent cell clusters in the presence of 10^9^ inhibitors. Approximately 100 Δ*pst* cells were plated on TG2PP and immediately incubated at 37°C. Top-agar carrying 10^9^ *phoA* cells was poured over the Δ*pst* bacteria at time 0, 3, 6, 12 or 24 h following Δ*pst* plating. The plates were then further incubated, such that the total incubation time for each plate was 48 h. The CFU number was counted at the end of the 48 h incubation. Each bar represents the mean ± S.E.M. of 5 independent cultures.

The selection of PCMs depends on the constitutive expression of AP, which hydrolyzes G2P releasing glycerol in sufficient amounts to allow bacterial growth at a reasonable rate. However, free glycerol may diffuse out of the periplasm of PCM cells and be captured by the numerous wild-type bacteria surrounding them, reducing glycerol concentration available for the growth of the PCM. If this sequence of events is correct, knockouts of genes related to glycerol metabolism in the ancestral inhibitor strain would diminish or abolish the growth inhibition of PCMs. To test this hypothesis, growth inhibition assays were performed with null mutants in genes associated with glycerol uptake and metabolism. The following gene deletions from the Keio collection were used in inhibition assays of PCM cells: *cyaA, crp, crr, glpA, glpB, glpC, glpD, glpF, glpK, glpQ, glpR, glpT, glpX*. *cyaA* and *crp* encode the CRP-cAMP transcriptional regulator and the bacterial adenylate cyclase, respectively. *crr* codes for the enzyme IIA^Glc^, which, among other things, activates adenylate cyclase [57, 58, 59]. *glpA, glpB* and *glpC* form an operon that encode a glycerol-3-phosphate dehydrogenase complex that converts glycerol-3-phosphate to dihydroxyacetone phosphate under anaerobic conditions [60] and *glpD* codes for an aerobic glycerol-3-phosphate dehydrogenase [61]. The *glpFKX* operon encodes the glycerol facilitator GlpF and the glycerol kinase GlpK, both directly involved in the uptake of glycerol [62], while GlpX is a fructose 1,6-bisphosphatase [63]. GlpR is the repressor of the glycerol-3-phosphate regulon. Noteworthy, some variants of strain MG1655, including the one originally sequenced [64] carry a 1 base-pair deletion in *glpR*, though our MG1655 variant harbors a wild-type *glpR* gene [65]. Finally, *glpQ* and *glpT* code for proteins involved in glycerol-3-phosphate uptake and metabolism [66]. 10^9^ bacteria each carrying single deletions of these genes were each mixed with 100 BW25113 Δ*pst* bacteria and plated on TG2PP plates (BW25113 is the parental strain of the Keio knockouts). The growth of Δ*pst* colonies was followed for 48 hours. Figure 3a shows that the *glpK* and *glpD* knockouts did not inhibit the growth of Δ*pst* colonies, while *crp, crr, cyaA, glpF* and *glpT* deletions inhibit only partially.

With the exception of *glpT*, a common characteristic of the knockout mutants that did not completely inhibited Δ*pst* growth is that none of them is able to grow with glycerol as the sole C source [62]. Herein one can conclude that bacteria that are unable to take up or metabolize glycerol do not inhibit the growth of PCMs. This by itself suggests that PCMs growth inhibition by the wild-type cells occurs because the glycerol produced in the periplasm of the PCM cells diffuses out and is predominantly consumed by the overwhelming excess of neighboring wild-type bacteria. In fact, the average frequency of PCMs is 5.0 · 10^−6^ (see Table 2), i. e., for each PCM cell there are ~ 20,000 competitor wild-type bacteria on the selective plate.

To test whether other individual genes might affect PCMs growth inhibition, the entire Keio library of mutants was screened. Figure 3b shows a picture of a typical experiment in which the mutants of Keio’s plate no. 53 were screened for their effect on Δ*pst* growth. In addition to the above mentioned genes, the only other knockout that is not auxotrophic and that did not inhibit the growth of Δ*pst* was *clpP* (not shown). The reason why the *clpP* knockout does not inhibit PCMs growth is unclear. All gene deletions that did not inhibit BW25113 Δ*pst* growth were transferred to strain MG1655 and tested again for growth inhibition of MG1655 Δ*pst* (Figure S3).

### Fitness of PCMs

The present hypothesis is that growth of most PCMs is inhibited by the surrounding wild-type cells due to competition for the limited glycerol. It may also suggests that the PCMs are less efficient in competing with the wild-type strain even though the hydrolysis of G2P and subsequent glycerol release occur in the PCM periplasm. To test this assumption we measured the fitness of the Δ*pst* mutant in the presence the wild-type strain with glycerol as the sole carbon source. Approximately 1000 bacteria of each strain - MG1655 and Δ*pst* (strain TC02) were mixed in medium TGlyP (0.2% glycerol as C source) and grown for 72 h with agitation. Samples were withdrawn every 24 hours, diluted and plated on Lagar supplemented with XP to differentiate between the strains. The selection coefficient of strain Δ*pst* at the end of 3 days of incubation was −0.67 ± 0.003, an average of three independent experiments. This suggests that in addition to the fact that the PCMs are vastly outnumber by the wild-type cells on the selective plate, they are also considerably less fit than the wild-type strain (by 67% per generation) when glycerol is used as a C source.

### How do some PCMs manage to grow after all?

Of the expected preexisting ~50,000 PCMs bacteria that are present among 10^9^ wild-type cells plated on selective TG2PP medium only a few dozens PCMs grow and eventually form colonies. However, there is not any impediment, in principle, for the growth of tens of thousands of PCM colonies in 48 h as shown in Figure S2. How then do some PCMs manage to escape the inhibition? Is it a stochastic phenomenon or do some lucky mutants have a unique characteristic that permit their growth? We suggest that some PCMs manage to grow and form visible colonies in cases that they randomly form upon plating occasional clusters of two or more cells. In these micro-clusters the glycerol released by one mutant is taken up by its adjacent PCM cell, which by mutual feeding allows further replication and the subsequent formation of a PCM colony. To test this hypothesis one hundred Δ*pst* cells were plated on each TG2PP plates. A thin layer of soft-agar containing 10^9^ Δ*phoA* cells was added at times 0, 3, 6, 12 and 24 hours on top of the plated Δ*pst* bacteria and the plates were further incubated for up to 48 h. Figure 3c shows that when 10^9^ *phoA* bacteria were added at time 0 h, no colony was visible after 48 h. However, when the *phoA* cells were added 3 hours later, around 12 colonies could be observed and in the 6 hours plate and over almost all Δ*pst* mutants managed to form colonies. Since the bacteria doubling time on TG2PP is around 2.5 hours, the 12 PCM colonies that grew after 3 hours could have arisen from clusters of 2 cooperating first-generation siblings. At 6 hours, when these clusters consisted of 4-8 siblings nearly all plated PCMs survived. This suggests that even small clusters of PCMs are able to grow and form colonies. Upon plating an overnight MG1655 culture containing ~ 50, 000 PCMs within a population of ~ 10^9^ wild-type cells, the probability for the formation of clusters consisting of different PCMs that would ensure growth on the selective plate is very small. Alternatively, a cluster of two or more cells may also originate from a single cell at a late phase of division (see Discussion).

### A mathematical model for PCMs growth and extinction

A mathematical model based on the tragedy of the commons exemplified by the competition for glycerol produced and released by the PCMs is detailed in the Appendix. The main hypothesis is that of spatial homogeneity, so that the model can be applied to the entire plate, or to a small portion of it (in this case assuming an absence of any sort of interaction with the surrounding adjacent regions). In addition to spatial homogeneity, the model assumes that both PCMs and wild-type cells compete equally for the glycerol moieties released by the mutant cells. The competition will eventually result in the extinction of both strains following G2P/glycerol depletion.

An ideal plate is considered in order to better understand the dynamics of this interaction. The main difference between an ideal plate and the actual TG2PP plate is that in the former the concentration of substrate is constant, while in the actual plate the concentration of G2P decreases with time. Analysis of the model reveals that both strains go to extinction unless they have the same fitness. In this case they can coexist provided there is a minimal initial number of mutants on the plate (see Equation 7 in the Appendix). In addition, PCMs may survive alone on the plate if the wild-type strain ability to capture glycerol is very low (see Equation 14). However, the results of the competition (see page 17) showed that the wild-type strain is considerably fitter than the PCM in the presence of glycerol as the sole carbon source (Equation 14 is not satisfied), suggesting that both strains go to extinction in the TG2PP plate. In fact, our data showed that the vast majority of the PCMs would never grow and form colonies and since the wild-type strain cannot grow on G2P both strains would eventually die.

As a numerical example for the model described in Equation 3, we may apply the parameters given in Table 2 of the Appendix. Assuming the carrying capacity *K* = 1, 000 and a starting number of 900 wild-type cells on a TG2PP plate with various amounts of PCMs we apply the system of equations described in Equation 3 taking into account the initial conditions *G*(0) = 0, *M*(0) = *M*_0_ and *W*(0) = *W*_0_ = 900, where *G*(0) is the initial G2P concentration, *W*(0) is the initial number of wild-type cells and *M*_0_ is the initial number of PCMs. If *M*_0_ = 10.2, both strains go to extinction, while for *M*_0_ = 10.3, they can coexist. Hence, both strains grow if the ratio between PCMs and wild-type cells is 1 : 89 (the higher the value of *M*_0_, the higher the level of glycerol produced by the PCMs, increasing the chances of coexistence). Naturally, the required ratio between PCMs and wild-type cells that allows coexistence would depend on the other values of the model. For instance, by increasing the G2P dissociation rate by 100%, that is, *α* = 0.2, the required PCMs:wild-type cells ratio becomes 1 : 175. Also, if the concentration of the surrounding wild-type bacteria increases, a higher number of PCMs is needed to form a colony. Since the real proportion of PCM to wild-type cells is around 1:20,000 (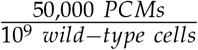, see Table 2), the fate of the vast majority of PCMs and wild-type cells is extinction.

## Discussion

Examples of bacterial growth inhibition by contact or proximity have been reported [54, 55, 56]. In one case, the inhibition was caused by toxins encoded by the CdiAB system from strain EC93, isolated from rat intestine [54]. Another instance was the inhibition caused by strains 25 and 256 isolated from cattle, whose mechanism of inhibition remains unknown [56]. In contrast to the aforementioned systems, stationary phase dependent contact inhibition reported by Lemonnier *et al.* [55] is the one that mostly resembles PCMs inhibition. It is the only one that occurs between isogenic strains, such that apparently the only difference between the inhibitor and inhibited strain is that the former carries a mutation in *glgC* that encodes an enzyme involved in glycogen synthesis. Bacteria carrying a mutation in this gene overproduce glycogen. However, the strain used in our study (MG1655) is not inhibited by the *glgC* mutant [55].

Krasovec *et al.* [22] have shown that high cell density partially inhibits the frequency of Rif^R^ mutants on a selective plate via a cell-cell *luxS*-dependent mechanism. A 77% reduction in cell density resulted in double mutation rate. PCMs mutation rate would probably be influenced by this cell density effect, but it only accounts for a small proportion of the observed eight hundred-fold inhibition.

In contrast to our previous view that the emergence of PCMs follows the pattern of stress-induced adaptive mutagenesis [26] it turned out that preexisting PCMs do exist and are inhibited because the overwhelming ancestral neighboring cells seize upon the glycerol moieties produced by the PCMs. Only bacterial strains that are unable to grow on glycerol do not inhibit PCMs growth (Figure 3a).

The data shown here strongly suggests that the late appearance of PCMs on the selective plate and their relatively low frequency is due to growth inhibition of the mutants. We cannot overrule, however, that some of these mutations occurred on the TG2PP selective plate and were induced by the medium conditions, as in the case of stress-induced adaptive mutations. Adaptive mutations occur in non-dividing bacteria and in the presence of scavenger cells [67]. In fact, we verified that the wild-type strain does not grow even for a few generations on the TG2PP plate (not shown). However, given the high number of preexisting PCMs in the culture and the extent of their inhibition, it is unlikely that any of the observed colonies of PCMs has derived from an adaptive mutation.

Our results have shown that mutagenesis under selective conditions may significantly mislead the estimate of the actual mutation rate. In the case of PCMs selection, from an initial rate of less than 2 × 10^−8^/gene/generation (assuming 100 mutants per 10^9^ bacteria and five *pst* genes as targets) the actual mutation rate (1.6 × 10^−6^/gene/generation) turned out to be close to the expected mutation rate in *E. coli* (~ 10^−6^/gene/generation [36, 68]).

PCMs inhibition by their ancestors is best represented as a “tragedy of the commons” case [33]. The Prisoner’s dilemma involves two players, which could cooperate or defect. Cooperation usually results in the greatest advantage for the majority of players, but defection benefits only the cheater [69]. In the “tragedy of the commons” both cheater (wild-type cells) and cooperator (PCM cells) loose because the public good (glycerol) is not produced sufficiently fast to provide a usable carbon source for the entire population or even to a significant part of the population of both cheaters or cooperators. In addition, the difference in fitness between the wild-type and PCMs strengthens the cheater and further magnifies the effect of the “tragedy”.

AP comprises up to six percent of total protein in wild-type bacteria under Pi-starvation [70]. We calculated the number of AP moieties present in a PCM bacterium by comparing the AP activity of Δ*pst* to the activity of known quantities of purified AP (Figure S4). Our data shows that there are on the average 2.67 · 10^−11^ *mg*/*ml* AP per bacterium. Assuming a MW of 94 kDa per AP homodimer, each bacterium harbors approximately 171,000 AP molecules. Given that an *E. coli* cell contains approximately 2,360,000 proteins [71], AP corresponds to 7.2% of Δ*pst*’s total protein. It is known that 0.2% of sugar is required to reach a yield of an OD_600_ = 2.0 (roughly 2 · 10^9^ bacteria/ml) [71]. In the case of G2P, whose carbon content is about 54% of the total molecule mass and in terms of *per capita* requirement, approximately 7.2 · 10^9^ molecules of G2P are consumed per bacterium. Thus, each AP unit should hydrolyze about 4.2 · 10^4^ G2P molecules to yield a single new bacterium.

The question of how some PCMs manage to grow could best be answered by assuming that the lucky colonies derive from small clusters of PCM cells. These clusters could be composed of different or identical PCM. In the former case, clumps of PCM cells would randomly repose on the selective plate upon plating and in the second case, the cluster would be composed of a bacterium undergoing replication. The evidence presented here (Figure 3c) supports the formation of homogeneous colonies, but it does not rule out the possibility that heterogeneous colonies formed by different PCM cells do occur. Moreover, these two alternatives are not mutually exclusive. In fact, a back of the envelope calculation shows that the probability that a pair of 2 different PCM cells would repose next to each other on the selective plate is about 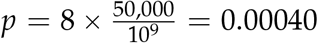, where 8 is an approximation for the number of immediate neighbor cells of each bacterium, 50, 000 is the estimated frequency of PCMs in a population of 10^9^ bacteria (based on Table 2). Thus in a population of 50,000 PCMs 20 PCM adjacent pairs are expected and a negligible number of 3-cell clusters. This number is in the range of those observed on the selective plates (see Figure 1). However, PCM clusters could also be homogeneous, i.e., composed of 2 siblings of a PCM cell that landed on the G2PP plate at some advanced step of division. All is needed is that some PCM cells would reproduce for 1-2 cycles on the selective plate using available traces of carbon sources.

In conclusion, the selection of PCMs on G2P is an example of how a normal mutant frequency could be hindered by the competition with their wild-type ancestors. This is also a case where cooperators (PCMs) are strongly inhibited by the overwhelming cheaters (wild-type cells), leading to a situation in which neither the cheaters nor the (majority) of cooperators can grow resulting in the “tragedy of the commons”.

## Methods

### Strains, plasmids and growth media

The bacterial strains used in this study are listed in Table S1. Unless otherwise stated, bacteria were incubated at 37°C under aerobic conditions. LB/L-agar is the standard rich medium [72]. The minimal media TGP and TG2PP were composed of T-salts [27] (0.12 M Tris-HCl, 80 mM NaCl, 20 mM KCl, 20 mM NH_4_Cl, 0.98 mM MgCl_2_ · 6 H_2_O, 2.46 mM Na_2_SO_4_, 2 mM CaCl_2_, 2 μM FeCl_3_, 2 μM ZnCl_2_, pH 7.5) supplemented with 1 mM of the KH_2_PO_4_ and either 0.2% glucose (TGP), 0.2% glycerol-2-phosphate (TG2PP) or 0.2% glycerol (TGlyP).

### Gene Knockouts

Genes and operons were deleted using the λ-*red* recombinase system as originally described by [73] and [74]. Briefly, chloramphenicol or kanamycin resistance genes were amplified using plasmid pKD3 or pKD4 as templates and the hybrid primers described below. The amplicons containing the *cat* gene or *kan* genes and 40 bases flanking sequences corresponding to genes *mutS* gene (primers mutS_mut_Fow – ATCACACCCCATTTAATATCAGGGAAC-CGGACATAACCCCGTGTAGGCTGGAGCTGCTTC and mutS_mut_Rev – GTTAATATTCCCGATAGCAAAAGACTATCGGGAATTGTTACATAT-GAATATCCTCCTTAG) and the operon *pstSCAB-phoU* (primers pst_m_Fow – GTCTGGT-GAATTATTTGTCGCTATCTTTCCCCGCCAGCAGTGTGTAGGCTGGAGCTGCTTC and pst_m_rev – AGGAGACATTATGAAAGTTATGCGTACCACCGTCGCAACTCATATGAATATCCTCCT-TAG) were electrotransformed into exponentially growing cultures of strains KM32 or KM44 grown in LB medium supplemented with 1 mM IPTG. Recombinants were selected on L-agar plates containing appropriate antibiotics. Knockout of *mutS* was confirmed by PCR with the primers mutS_ver_Fow and mutS_ver_Rev and the deletion of *pstSCAB-phoU* was confirmed by PCR and AP assay. When required, gene deletions were transferred to strain MG1655 by P1 transduction.

### P1 transduction

Chromosomal markers were transferred between strains using P1 transduction as described in Miller, 1992 [72]. Mutants carrying specific deletions were selected on L-agar plates supplemented with the appropriate antibiotic. Further confirmation of the genetic marker was performed by PCR. Following transduction of gene deletions, the antibiotic marker was removed in some instances using plasmid pFLP2 as described [73].

### Selection of PCMs on TG2PP medium

PCMs were selected as described [25, 26]. Briefly, bacteria were grown overnight in TGP medium, washed three times with 0.9% NaCl and plated on TG2PP supplemented with XP (40 μ g/ml). Approximately 10^9^ were plated on each plate, incubated for different time lengths and counted daily, as specified in the text. Blue colonies were considered PCMs.

### PCMs inhibition assay

Δ*pst* cells and the inhibitor strain to be tested were grown overnight in TGP medium. On the next day, both cultures were washed three times with 0.9% NaCl and usually 10^9^ inhibitor cells and 100 Δ*pst* bacteria were mixed and plated on TG2PP medium. Plates were incubated for 2 days and the emergence of blue colonies was recorded.

### Survival of wild-type cells on TG2PP plates

Wild-type bacteria were plated on TG2PP as described above. Agar plugs with 5 mm diameter were removed every 24 h from the plate with the help of a glass cannula. Bacteria were eluted from the agar plug by vortexing for 1 minute. Bacteria were then diluted and plated on L-agar plates.

### Bacterial lysis by freezing/thawing

Bacteria were grown overnight in TGP medium and concentrated to a final concentration of 10^10^ bacteria/ml. An aliquot of 100 μl was centrifuged and the bacteria were resuspended in the same volume of lysis solution (1 M Tris pH 8,0, 5 mg/ml lysozyme, 100 mM phenyl-methylsulfonyl fluoride and 1 μl DNase). The suspension was kept on ice for 30 minutes and then submitted to 6 cycles of freezing/thawing by immersion in liquid nitrogen. The lysed bacteria were centrifuged to precipitate the debris and the supernatant was kept on ice until further use.

### Fluctuation test

Strains MG1655 and RI103 (MG1655 Δ*mutS*∷Cm) were grown overnight in medium TGP. On the next day, the cultures were washed three times in 0.9% NaCl. Approximately one thousand bacteria from each culture were inoculated in each of 60 wells filled with 100 μl TGP medium containing 110 nM glucose and incubated at 37°C for 30 hours. The total volume of the cultures was then plated on TG2PP plates which were incubated for 48 hours, when the frequency of PCMs was assessed. The number of bacteria in each culture (total CFU) was counted in parallel cultures. To calculate the mutation rate of PCMs derived from MG1655 the P_0_ method was employed [7]. The mutation rate of strain RI103 was calculated using the P_0_ method and also by the Lea & Coulson [48] and Jones [49] methods.

### Competition assay

MG1655 and the Δ*pst* mutant (strain TC02) were grown overnight in medium TGP. On the next morning, approximately 1000 cells of each strain were mixed in 5 ml of medium TGlyP and grown at 37°Cwith agitation. Samples were taken at 0, 24 and 48 h, diluted and plated on L-agar supplemented with XP. The selection coefficient was calculated from the growth rates according to the formula: 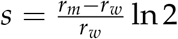, where *r_m_* and *r_w_* represent the growth rate of the mutant and wild-type strain, respectively. The competition was performed in triplicates.

### Growth inhibition of PCM clusters

Overnight cultures of MG1655 Δ*pst* (strain TC02) grown in TGP medium were washed and diluted in 0.9% NaCl. Approximately one hundred bacteria were then plated on TG2PP. The plates were incubated at 37°C for 0, 3, 6, 12 or 24 hours, at which times a layer of soft-agar (6 g/l) containing 10^9^ Δ*phoA* cells was poured over the plate. The plates were returned to the incubator until the total time of 48 hours and the number of PCM colonies was counted.

### Screening of the Keio collection

The Keio strains and the Δ*pst* mutant (strain TC01) were grown overnight in TGP medium. On the next day, 1 ml of each culture was centrifuged and resuspended in the same volume of 0.9% NaCl. The Δ*pst* suspension was diluted 100X and 30 μl of this dilution were used to create each of the linear patches of bacteria on a TG2PP plate. Once the Δ*pst* patch was dry, 2 μl of the each Keio strain were dropped over the patch. The plates were incubated for 48 hours at 37°C and the growth of Δ*pst* inside the circle formed by the Keio strain sample was evaluated. Growth inhibition was characterized by an empty white circle while a circle filled with a blue patch indicated that the Keio strain did not inhibit the growth of Δ*pst*. The Keio knockouts that did not inhibit Δ*pst* growth were further tested in a conventional inhibition assay to confirm this phenotype.

### Assessment of Rif^R^ mutant frequency

Bacteria (strain MG1655) grown overnight in medium TGP were washed three times with 0.9% NaCl. Approximately 10^9^ bacteria were plated on TGP plates containing 100 μg/ml rifampicin. Mutant frequency was calculated as the ratio of resistant mutants over the total number of plated bacteria, estimated by CFU counting of bacteria on L-agar plates.

### PCMs growth Inhibition through filters

Bacteria (strains TC01(Δ*pst*) and RI05 (Δ*phoA*)) were grown overnight in TGP medium. On the next day cultures were washed three times with 0.9% NaCl and 10^9^ Δ*phoA* cells were spread on a TG2PP plate. A sterile acetate cellulose filter or a stack of 3 filters (pore size μm) were placed at the center of the plate and a drop containing approximately 100 Δ*pst* cells was applied on the surface of the filter. Plates were incubated for 48 hours and the growth of PCM colonies was evaluated.

## Acknowledgements

We are grateful to Fundação de Amparo à Pesquisa do Estado de São Paulo (FAPESP) for supporting this study. BS is a recipient of the Conselho Nacional de Desenvolvimento Científico e Tecnológico (CNPq) productivity scholarship. HIN, GTM and TCSR were supported by FAPESP scholarships. We also thanks CAPES (Coordenação de Aperfeiçoamento de Pessoal de Nível Superior) for partially supporting this study.

## References

[1] Ferenci TT, Maharjan R. A shifting mutational landscape in 6 nutritional states: Stress-induced mutagenesis as a series of distinct stress inputmutation output relationships. PLOS Biology. 2017;.

[2] Ferenci T. Irregularities in genetic variation and mutation rates with environmental stresses. Environmental microbiology. 2019;.

[3] Al Mamun AAM, Lombardo MJ, Shee C, Lisewski AM, Gonzalez C, Lin D, et al. Identity and function of a large gene network underlying mutagenic repair of DNA breaks. Science. 2012;338(6112):1344–1348.

[4] Foster PL. Stress-induced mutagenesis in bacteria. Crit Rev Biochem Mol Biol. 2007;42(5):373–397. Available from: http://dx.doi.org/10.1080/10409230701648494.

[5] Galhardo RS, Hastings PJ, Rosenberg SM. Mutation as a stress response and the regulation of evolvability. Crit Rev Biochem Mol Biol. 2007;42(5):399–435. Available from: http://dx.doi.org/10.1080/10409230701648502.

[6] Maharjan RP, Ferenci T. A shifting mutational landscape in 6 nutritional states: Stress-induced mutagenesis as a series of distinct stress input-mutation output relationships. PLoS biology. 2017 Jun;15:e2001477.

[7] Luria S, Delbruck M. Mutations of bacteria from virus sensitivity to virus resistance. Genetics. 1943;28(6):491–511.

[8] Cairns J, Overbaugh J, Miller S. The origin of mutants. Nature. 1988 Sep;335(6186):142–145. Available from: http://dx.doi.org/10.1038/335142a0.

[9] Hall BG. Adaptive mutations in *Escherichia coli* as a model for the multiple mutational origins of tumors. Proceedings of the National Academy of Sciences. 1995;92(12):5669–5673.

[10] Cohen SE, Walker GC. The transcription elongation factor NusA is required for stress-induced mutagenesis in *Escherichia coli*. Curr Biol. 2010 Jan;20(1):80–85. Available from: http://dx.doi.org/10.1016/j.cub.2009.11.039.

[11] Fitzgerald DM, Rosenberg SM. What is mutation? A chapter in the series: How microbes “jeopardize” the modern synthesis. PLoS genetics. 2019;15(4):e1007995.

[12] Roth JR, Kugelberg E, Reams AB, Kofoid E, Andersson DI. Origin of mutations under selection: the adaptive mutation controversy. Annu Rev Microbiol. 2006;60:477–501. Available from: http://dx.doi.org/10.1146/annurev.micro.60.080805.142045.

[13] Maisnier-Patin S, Roth J. The origin of mutants under selection: how natural selection mimics mutagenesis (adaptive mutation). Cold Spring Harb Perspect Biol 7: a018176. Cold Spring Harbor perspectives in biology. 2015;.

[14] Yosef I, Edgar R, Levy A, Amitai G, Sorek R, Munitz A, et al. Natural selection underlies apparent stress-induced mutagenesis in a bacteriophage infection model. Nature microbiology. 2016;1(6):16047.

[15] Zhang Z, Saier Jr MH. Transposon-mediated activation of the *Escherichia coli glpFK* operon is inhibited by specific DNA-binding proteins: Implications for stress-induced transposition events. Mutation Research/Fundamental and Molecular Mechanisms of Mutagenesis. 2016;793:22–31.

[16] Foster PL. Rebuttal: growth under selection stimulates Lac(+) reversion (Roth and Andersson). Journal of bacteriology. 2004 Aug;186:4861.

[17] Cairns J, Foster PL. Adaptive reversion of a frameshift mutation in *Escherichia coli*. Genetics. 1991;128(4):695–701.

[18] Hoffmann GR, Gray CL, Lange PB, Marando CI. A source of artifact in the *lacZ* reversion assay in *Escherichia coli*. Mutation research Genetic toxicology and environmental mutagenesis. 2015 Jun;784-785:23–30.

[19] Smith TF, Sadler JR. The nature of lactose operator constitutive mutations. Journal of molecular biology. 1971;59(2):273–305.

[20] Tang M, Wang L, Zhang P, Huang P, Wu L. Use Pgal agarose minimal plate to screen *lac* constitutive mutation. Annals of microbiology. 2009;59(1):179.

[21] Miller JH, Funchain P, Clendenin W, Huang T, Nguyen A, Wolff E, et al. *Escherichia coli* strains (ndk) lacking nucleoside diphosphate kinase are powerful mutators for base substitutions and frameshifts in mismatch-repair-deficient strains. Genetics. 2002 Sep;162:5–13.

[22] Krašovec R, Belavkin RV, Aston JAD, Channon A, Aston E, Rash BM, et al. Mutation rate plasticity in rifampicin resistance depends on *Escherichia coli* cell-cell interactions. Nature communications. 2014 Apr;5:3742.

[23] Richards H, Gifford DR, Hatcher C, Faulkner KJ, Belavkin RV, Channon A, et al. Spontaneous mutation rate is a plastic trait associated with population density across domains of life. PLoS biology. 2017;15(8):e2002731.

[24] Gardner S, McCleary W. Control of the phoBR Regulon in *Escherichia coli*. EcoSal Plus. 2019;8(2).

[25] Torriani A, Rothman F. Mutants of *Escherichia coli* constitutive for alkaline phosphatase [Note]. J Bacteriol. 1961;81(5):835–836.

[26] Neves HI, Pereira TF, Yagil E, Spira B. Ugp and PitA participate in the selection of PHO-constitutive mutants. Journal of bacteriology. 2015;197(8):1378–1385.

[27] Echols H, Garen A, Garen S, Torriani A. Genetic control of repression of alkaline phosphatase in *E. coli*. J Mol Biol. 1961 Aug;3:425–38.

[28] Aguena M, Spira B. Transcriptional processing of the *pst* operon of *Escherichia coli*. Current microbiology. 2009;58(3):264–267.

[29] Wanner BL. 87. Phosphorus assimilation and control of the phosphate regulon. In: Neidhardt F, Curtis R, Ingraham J, Lin E, Low K, Magasanik B, et al., editors. *Escherichia coli* and *Salmonella*: Cellular and Molecular Biology. 2nd ed. American Society for Microbiology; 1996. p. 1357–1381.

[30] Heller KB, Lin EC, Wilson TH. Substrate specificity and transport properties of the glycerol facilitator of *Escherichia coli*. J Bacteriol. 1980 Oct;144(1):274–278.

[31] Tong H, Hu Q, Zhu L, Dong X. Prokaryotic Aquaporins. Cells. 2019;8(11):1316.

[32] Yang K, Wang M, Metcalf WW. Uptake of glycerol-2-phosphate via the *ugp*-encoded transporter in *Escherichia coli* K-12. J Bacteriol. 2009 Jul;191(14):4667–4670. Available from: http://dx.doi.org/10.1128/JB.00235-09.

[33] Schuster M, Foxall E, Finch D, Smith H, De Leenheer P. Tragedy of the commons in the chemostat. PloS one. 2017;12(12):e0186119.

[34] Dandekar AA, Chugani S, Greenberg EP. Bacterial quorum sensing and metabolic incentives to cooperate. Science. 2012;338(6104):264–266.

[35] Rainey PB, Rainey K. Evolution of cooperation and conflict in experimental bacterial populations. Nature. 2003;425(6953):72.

[36] Drake JW, Charlesworth B, Charlesworth D, Crow JF. Rates of spontaneous mutation. Genetics. 1998 Apr;148:1667–1686.

[37] Lee H, Popodi E, Tang H, Foster PL. Rate and molecular spectrum of spontaneous mutations in the bacterium *Escherichia coli* as determined by whole-genome sequencing. Proceedings of the National Academy of Sciences. 2012;109(41):E2774–E2783.

[38] Maharjan RP, Ferenci T. *Escherichia coli* mutation rates and spectra with combinations of environmental limitations. Microbiology. 2018;164(12):1491–1502.

[39] Chueca B, Berdejo D, Gomes-Neto NJ, Pagán R, García-Gonzalo D. Emergence of Hyper-Resistant *Escherichia coli* MG1655 Derivative Strains after Applying Sub-Inhibitory Doses of Individual Constituents of Essential Oils. Frontiers in microbiology. 2016;7:273.

[40] Thi TD, López E, Rodríguez-Rojas A, Rodríguez-Beltrán J, Couce A, Guelfo JR, et al. Effect of *recA* inactivation on mutagenesis of *Escherichia coli* exposed to sublethal concentrations of antimicrobials. The Journal of antimicrobial chemotherapy. 2011 Mar;66:531–538.

[41] Baharoglu Z, Mazel D. Vibrio cholerae triggers SOS and mutagenesis in response to a wide range of antibiotics: a route towards multiresistance. Antimicrobial agents and chemotherapy. 2011 May;55:2438–2441.

[42] Elez M, Radman M, Matic I. The frequency and structure of recombinant products is determined by the cellular level of MutL. Proceedings of the National Academy of Sciences of the United States of America. 2007 May;104:8935–8940.

[43] Ibacache-Quiroga C, Oliveros JC, Couce A, Blázquez J. Parallel evolution of high-level aminoglycoside resistance in *Escherichia coli* under low and high mutation supply rates. Frontiers in microbiology. 2018;9:427.

[44] Cox EC, Degnen GE, Scheppe ML. Mutator gene studies in *Escherichia coli*: the *mutS* gene. Genetics. 1972;72(4):551–567.

[45] LeClerc JE, Li B, Payne WL, Cebula TA. High mutation frequencies among *Escherichia coli* and Salmonella pathogens. Science. 1996;274(5290):1208–1211.

[46] Li GM. Mechanisms and functions of DNA mismatch repair. Cell research. 2008 Jan;18:85–98.

[47] Foster PL. Methods for determining spontaneous mutation rates. Methods Enzymol. 2006;409:195–213. Available from: http://dx.doi.org/10.1016/S0076-6879(05)09012-9.

[48] Lea D, Coulson C. The distribution of the numbers of mutants in bacterial populations. Journal of genetics. 1949 Dec;49:264–285.

[49] Jones M. Accounting for plating efficiency when estimating spontaneous mutation rates. Mutation Research. 1993 Oct;292:187–189.

[50] Rosche WA, Foster PL. Determining mutation rates in bacterial populations. Methods (San Diego, Calif). 2000 Jan;20:4–17.

[51] Galan JC, Turrientes MC, Baquero MR, Rodriguez-Alcayna M, Martinez-Amado J, Martinez JL, et al. Mutation rate is reduced by increased dosage of mutL gene in Escherichia coli K-12. FEMS microbiology letters. 2007 Oct;275:263–269.

[52] Kolodkin-Gal I, Hazan R, Gaathon A, Carmeli S, Engelberg-Kulka H. A linear pentapeptide is a quorum-sensing factor required for *mazEF*-mediated cell death in *Escherichia coli*. science. 2007;318(5850):652–655.

[53] Aoki SK, Pamma R, Hernday AD, Bickham JE, Braaten BA, Low DA. Contact-dependent inhibition of growth in *Escherichia coli*. Science. 2005;309(5738):1245–1248.

[54] Aoki SK, Diner EJ, de Roodenbeke CT, Burgess BR, Poole SJ, Braaten BA, et al. A widespread family of polymorphic contact-dependent toxin delivery systems in bacteria. Nature. 2010 Nov;468:439–442.

[55] Lemonnier M, Levin BR, Romeo T, Garner K, Baquero MR, Mercante J, et al. The evolution of contact-dependent inhibition in non-growing populations of *Escherichia coli*. Proceedings Biological sciences. 2008 Jan;275:3–10.

[56] Sawant AA, Casavant NC, Call DR, Besser TE. Proximity-dependent inhibition in *Escherichia coli* isolates from cattle. Applied and environmental microbiology. 2011 Apr;77:2345–2351.

[57] Westfall CS, Levin PA. Comprehensive analysis of central carbon metabolism illuminates connections between nutrient availability, growth rate, and cell morphology in *Escherichia coli*. PLoS genetics. 2018 Feb;14:e1007205.

[58] Park YH, Lee BR, Seok YJ, Peterkofsky A. In vitro reconstitution of catabolite repression in *Escherichia coli*. The Journal of biological chemistry. 2006 Mar;281:6448–6454.

[59] Saier M, Feucht B, Hofstadter L. Regulation of carbohydrate uptake and adenylate cyclase activity mediated by the enzymes II of the phosphoenolpyruvate: sugar phospho-transferase system in *Escherichia coli*. Journal of Biological Chemistry. 1976;251(3):883–892.

[60] Cole S, Eiglmeier K, Ahmed S, Honore N, Elmes L, Anderson W, et al. Nucleotide sequence and gene-polypeptide relationships of the *glpABC* operon encoding the anaerobic sn-glycerol-3-phosphate dehydrogenase of *Escherichia coli* K-12. Journal of bacteriology. 1988;170(6):2448–2456.

[61] Yeh JI, Chinte U, Du S. Structure of glycerol-3-phosphate dehydrogenase, an essential monotopic membrane enzyme involved in respiration and metabolism. Proceedings of the National Academy of Sciences. 2008;105(9):3280–3285.

[62] Joyce AR, Reed JL, White A, Edwards R, Osterman A, Baba T, et al. Experimental and computational assessment of conditionally essential genes in *Escherichia coli*. Journal of bacteriology. 2006;188(23):8259–8271.

[63] Brown G, Singer A, Lunin VV, Proudfoot M, Skarina T, Flick R, et al. Structural and biochemical characterization of the type II fructose-1,6-bisphosphatase GlpX from *Escherichia coli*. The Journal of biological chemistry. 2009 Feb;284:3784–3792.

[64] Blattner FR, Plunkett G, Bloch CA, Perna NT, Burland V, Riley M, et al. The complete genome sequence of *Escherichia coli* K-12. Science (New York, NY). 1997 Sep;277:1453–1462.

[65] Stasi R, Neves HI, Spira B. Phosphate uptake by the phosphonate transport system PhnCDE. BMC microbiology. 2019 Apr;19:79.

[66] Larson TJ, Ehrmann M, Boos W. Periplasmic glycerophosphodiester phosphodiesterase of *Escherichia coli*, a new enzyme of the glp regulon. The Journal of biological chemistry. 1983 May;258:5428–5432.

[67] Williams AB, Foster PL. Stress-Induced Mutagenesis. EcoSal Plus. 2012 Nov;5.

[68] Whitehead DJ, Wilke CO, Vernazobres D, Bornberg-Bauer E. The look-ahead effect of phenotypic mutations. Biology Direct. 2008;3(1):18.

[69] Axelrod R. The evolution of Cooperation. New York: Basic Books; 1984.

[70] Garen A, C Levinthal C. A fine-structure genetic and chemical study of the enzyme alkaline phosphatase of *E. coli*. I. Purification and characterization of alkaline phosphatase. Biochim Biophys Acta. 1960 Mar;38:470–483.

[71] Milo R, Jorgensen P, Moran U, Weber G, Springer M. BioNumbers-the database of key numbers in molecular and cell biology. Nucleic acids research. 2009;38(suppl_1):D750–D753.

[72] Miller JH. A Short Course In Bacterial Genetics: A Laboratory Manual And Handbook For *Escherichia coli* And Related Bacteria. Cold Spring Harbor Laboratory, Cold Spring Harbor, N.Y.; 1992.

[73] Datsenko KA, Wanner BL. One-step inactivation of chromosomal genes in *Escherichia coli* K-12 using PCR products. Proc Natl Acad Sci USA. 2000 Jun;97(12):6640–5.

[74] Murphy KC, Campellone KG, Poteete AR. PCR-mediated gene replacement in *Escherichia coli*. Gene. 2000 Apr;246(1-2):321–330.

